## Supplementary Material for "Human orbitofrontal cortex signals decision outcomes to sensory cortex during flexible tactile learning"

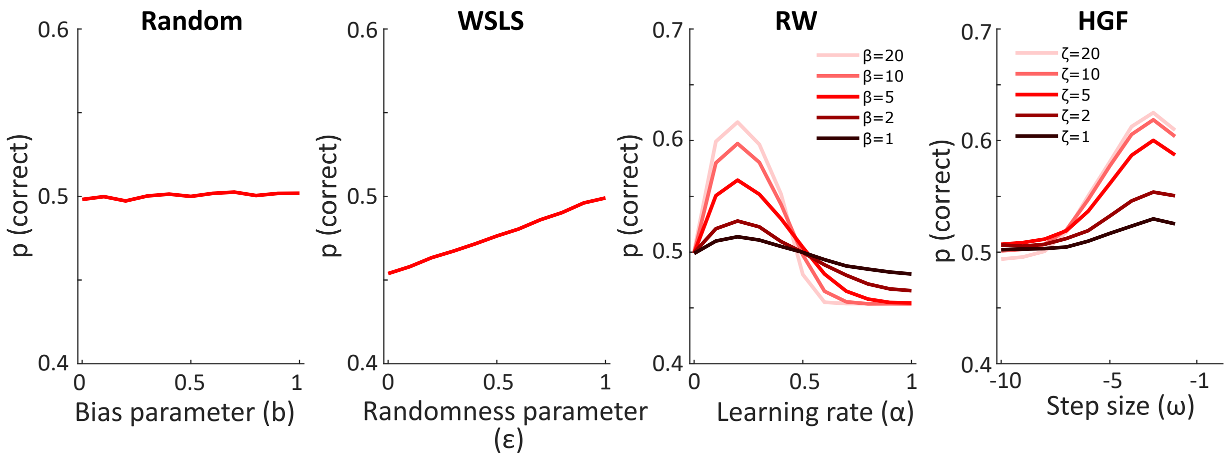


**Supplementary Fig. 1:** Four model simulations (500 per parameter setting) show how the free parameters influence behavior (Random Responding (Random), Win-Stay-Lose-Switch (WSLS), Rescorla-Wagner (RW) and Hierarchical Gaussian Filter (HGF)).

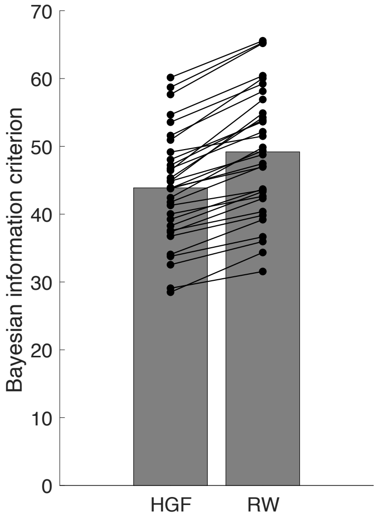


**Supplementary Fig. 2:** The Bayesian information criterion (BIC) for both HGF and RW model. The smaller the value, the better the model fit. Each dot represents a single participant.

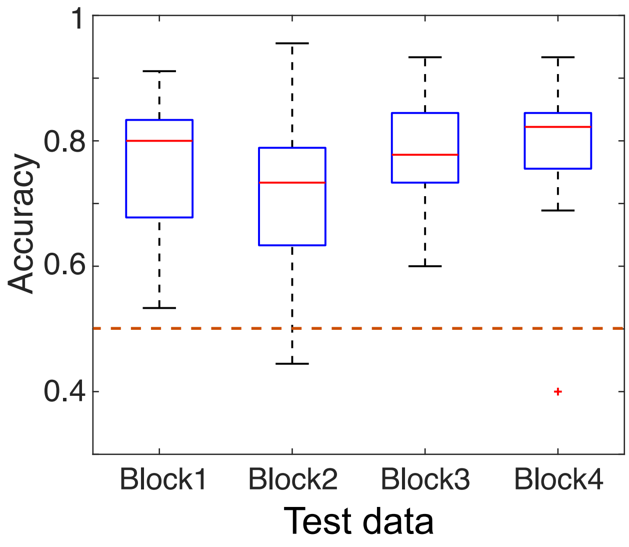


**Supplementary Fig. 3**: The cross-validation test for the HGF model. The parameters ($\omega$ and $\zeta$) of the HGF model were estimated by fitting to the real behavioral responses in the training data (i.e., run1 and run2). These parameters were then used to predict behavior in the test data (i.e., run3). The prediction accuracy was calculated, which revealed the generalizability of the model across different data samples. Box plots indicate median (middle line), 25th, 75th percentile (box) and the maximum and minimum (whiskers) as well as outlier (red cross). The dashed red line indicates chance level.

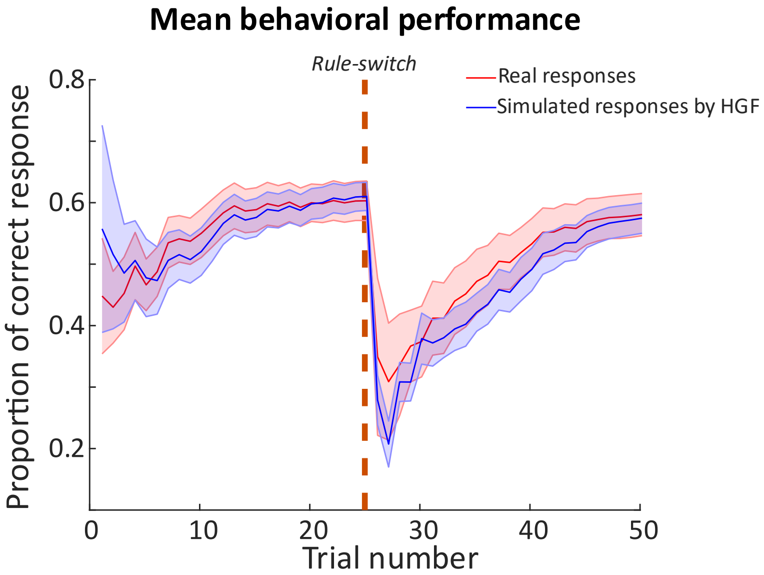


**Supplementary Fig. 4:** The overlay of real behavioral responses and the simulated responses produced by the winning HGF model using the optimized parameters from model fitting. The shaded area indicates the standard error of the mean (SEM).

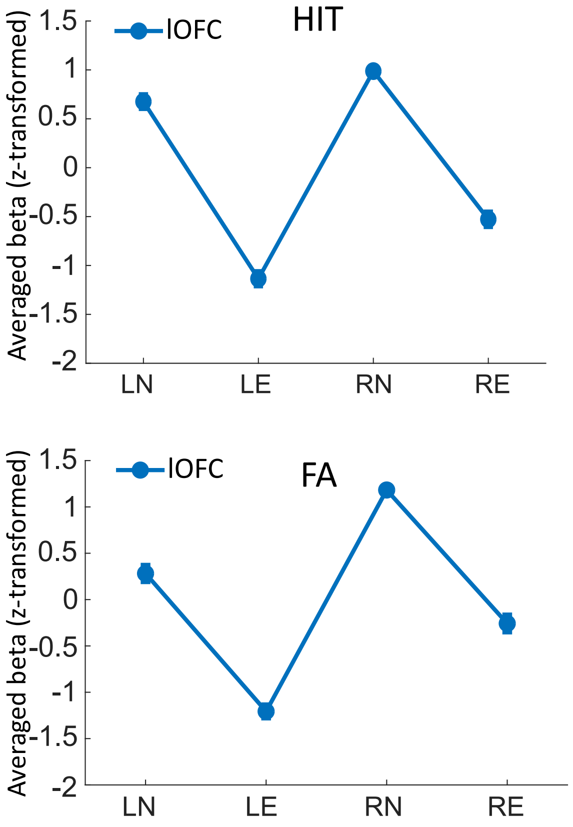


**Supplementary Fig. 5:** The BOLD signals in lOFC across the four key learning phases for the four types of trials (HIT, CR, FA, MISS) respectively. The error bars indicate the SEM.

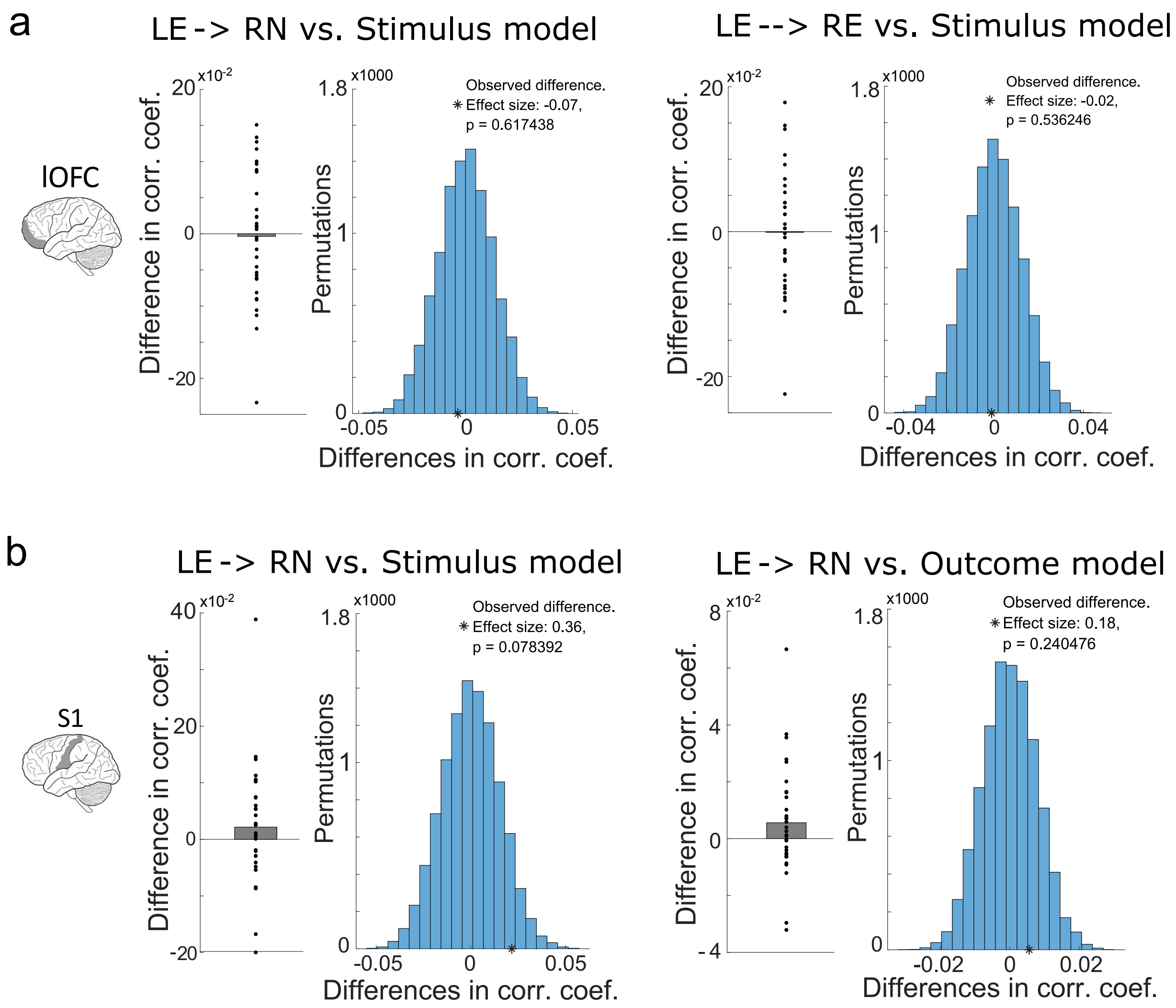


**Supplementary Fig. 6:** The non-significant results for RSA analysis. a. The response patterns in lOFC did not significantly represent the stimulus during both the immediate effect of the reversal (LE->RN) and the stable adaptation after re-learning (LE-->RE). b. The response patterns in S1 did not significantly represent both the stimulus and outcome during the immediate effect of the reversal (LE->RN). For both lOFC and S1, the group mean was compared using both one-sided Wilcoxon signed-rank test (left) and permutation test (right).

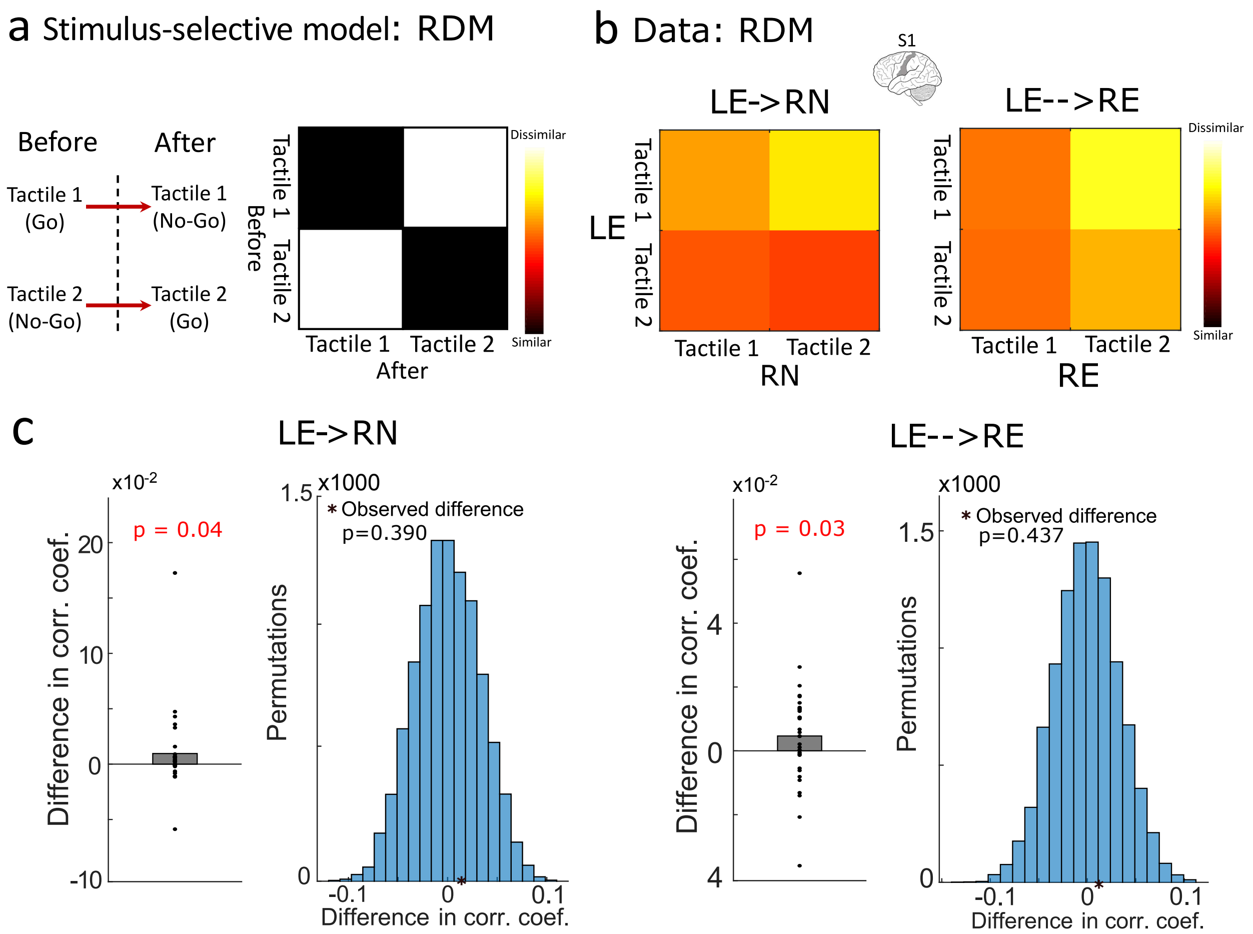


**Supplementary Fig. 7:** The RSA for stimulus-selectivity in S1, time-locked to stimulus presentation. a. Description of the stimulus-selective model and the corresponding representational dissimilarity matrix (RDM). b. The RDM for stimulus-selective responses from S1 for both immediately after reversals (LE->RN) and during re-leaning (LE-->RE). c. The comparison between the mean 'similar' (black elements in model RDMs) and mean 'dissimilar' (white elements in model RDMs) using a one-sided Wilcoxon signed-rank test together with permutation tests for LE->RN and LE-->RE.

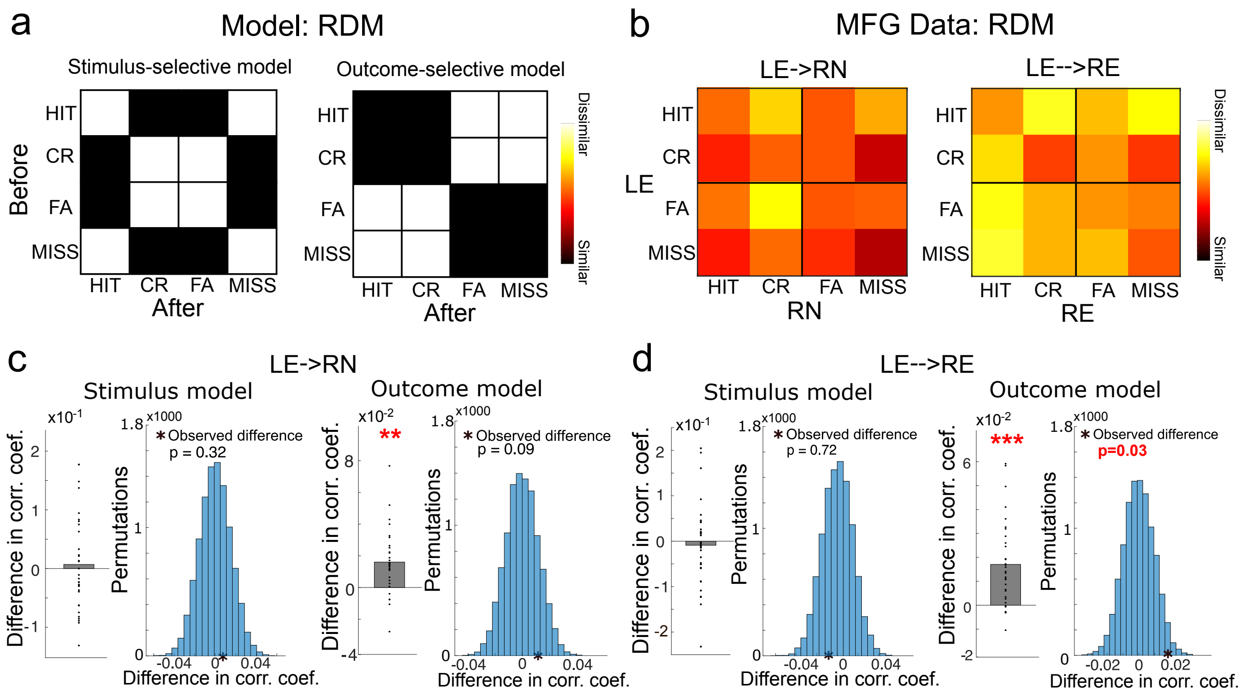


**Supplementary Fig. 8:** The RSA analyses for MFG. a. The representational dissimilarity matrix (RDM) of stimulus-selective and outcome-selective model. b. The RDM of response pattern in MFG for both immediately after reversals (LE->RN) and during re-leaning (LE-->RE). c. The comparison between the mean 'similar' (black elements in model RDMs) and mean 'dissimilar' (white elements in model RDMs) using a one-sided Wilcoxon signed-rank test together with permutation tests after reversals (LE->RN) and d. same with c but during re-leaning (LE-->RE). ** indicates $p < 0.01$, *** indicates $p < 0.001$.

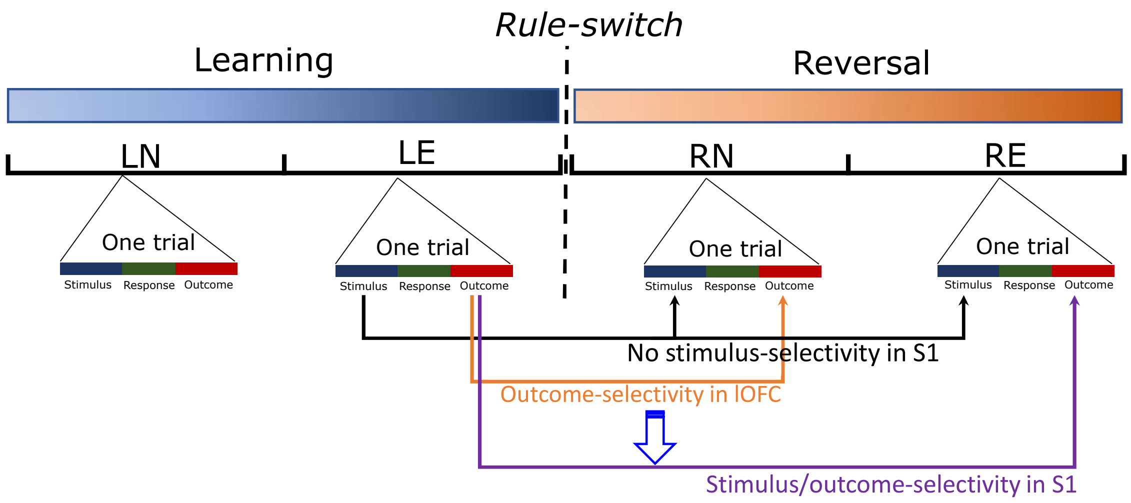


**Supplementary Fig. 9:** The schematic for the representation of stimulus and outcome in S1 and lOFC after reversal.

**Supplementary Table 1.** The Bayes optimal parameters for perceptual model in both RW and HGF

| | α in RW | ω in HGF | | α in RW | ω in HGF |
|---|---|---|---|---|---|
| Block1 | 0.143 | -3.25 | Block7 | 0.147 | -3.03 |
| Block2 | 0.137 | -3.61 | Block8 | 0.141 | -3.19 |
| Block3 | 0.141 | -3.33 | Block9 | 0.142 | -3.38 |
| Block4 | 0.150 | -2.88 | Block10 | 0.138 | -3.52 |
| Block5 | 0.140 | -3.65 | Block11 | 0.147 | -2.99 |
| Block6 | 0.143 | -3.04 | Block12 | 0.137 | -3.49 |

**Supplementary Table 2.** The maximum a posteriori estimates of the free parameters from model fitting (means and standard errors (SD)) for the RW and HGF model

| Models | Parameters | Mean | SD |
|---|---|---|---|
| RW | α | 0.19 | 0.03 |
| | $\beta$ | 5.68 | 3.13 |
| HGF | ω | -3.45 | 0.59 |
| | $\zeta$ | 13.48 | 2.59 |

**Supplementary Table 3.** Brain regions positively related to the outcome prediction error ($p < 0.001$, uncorrected)

| Regions | Hemisphere | Peak coordinates | | | T-score |
|---|---|---|---|---|---|
| | | x | y | z | |
| Insula | R | 30 | 22 | 2 | 5.57 |
| Insula | L | -28 | 22 | -2 | 5.82 |
| Posterior parietal cortex | R | 34 | -64 | 44 | 5.90 |
| Orbitofrontal cortex | R | 40 | 48 | -2 | 4.99 |
| Middle frontal gyrus (inf) | R | 46 | 14 | 32 | 6.77 |
| Middle frontal gyrus (sup) | R | 48 | 12 | 50 | 4.99 |
| Supplementary motor area | R&L | -4 | 26 | 42 | 5.31 |

We found two distinct clusters in middle frontal gyrus: “Inf” indicates its inferior part and “sup” indicates its superior part.
